# Model-based analysis of response and resistance factors of cetuximab treatment in gastric cancer cell lines

**DOI:** 10.1101/656967

**Authors:** Elba Raimúndez, Simone Keller, Gwen Zwingenberger, Karolin Ebert, Sabine Hug, Fabian J. Theis, Dieter Maier, Birgit Luber, Jan Hasenauer

**Affiliations:** Helmholtz Zentrum München-German Research Center for Environmental Health, Institute of Computational Biology, 85764 Neuherberg, Germany; Center for Mathematics, Technische Universität München, 85748 Garching, Germany; Technical University of Munich, School of Medicine, Klinikum rechts der Isar, Institute of Pathology, 81675 München, Germany; Biomax Informatics AG, Planegg; Faculty of Mathematics and Natural Sciences, University of Bonn, 53113 Bonn, Germany

## Abstract

Targeted cancer therapies are powerful alternatives to chemotherapies or can be used complementary to these. Yet, the response to targeted treatments depends on a variety of factors, including mutations and expression levels, and therefore their outcome is difficult to predict. Here, we develop a mechanistic model of gastric cancer to study response and resistance factors for cetuximab treatment. The model captures the EGFR, ERK and AKT signaling pathways in two gastric cancer cell lines with different mutation patterns. We train the model using a comprehensive selection of time and dose response measurements, and provide an assessment of parameter and prediction uncertainties. We demonstrate that the proposed model facilitates the identification of causal differences between the cell lines. Furthermore, our study shows that the model provides accurate predictions for the responses to different perturbations, such as knockdown and knockout experiments. Among other results, the model predicted the effect of MET mutations on cetuximab sensitivity. These predictive capabilities render the model a powerful basis for the assessment of gastric cancer signaling and for the development and discovery of predictive biomarkers.

**Author Summary:** Unraveling the causal differences between drug responders and non-responders is an important challenge. The information can help to understand molecular mechanisms and to guide the selection and design of targeted therapies. Here, we approach this problem for cetuximab treatment for gastric cancer using mechanistic mathematical modeling. The proposed model describes multiple gastric cancer cell lines and can accurately predict the response in several validation experiments. Our analysis provides a differentiated view on mutations and explains, for instance, the relevance of MET mutations and the insignificance of PIK3CA mutation in the considered cell lines. The model might provide the basis for understanding the recent failure of several clinical studies.

## 1 Background

Gastric cancer is the fifth most common cancer and third leading cause of death from cancer worldwide (Bray et al., 2018). Treatment options include surgery, chemo- and radiation therapy. However, the overall survival rate remains unsatisfactory due to molecular and clinical heterogeneity (Lordick et al., 2014), therefore new treatment options are urgently required. Novel drugs targeting members of a family of receptor tyrosine kinases including the epidermal growth factor receptor (EGFR) have shown mixed success in clinical trials (Bang et al., 2010; Lordick et al., 2013). Among others, the EGFR antibody cetuximab did not improve patient survival in a phase III clinical trial (Lordick et al., 2013).

A potential reason for the failure of cetuximab is the molecular heterogeneity of gastric cancer (Lordick et al., 2014). Due to this heterogeneity, only a small subgroup of patients might benefit from the targeted therapy. However, a suitable biomarker for patient stratification is currently not available. Here, we aim to understand response and resistance mechanisms for cetuximab treatment in gastric cancer, to unravel causal differences between cetuximab responders and non-responders (Keller et al., 2017), and to identify biomarkers for guiding targeted therapy by using a cell culture model.

Conceptually, biomarkers for responsive patient subgroups can be identified using statistical approaches characterizing responder and non-responder subgroups within different molecular high-throughput methods. Unfortunately, the necessary large-scale studies on the response of gastric cancer patients to cetuximab are missing. In addition, many proposed biomarkers from purely associative approaches have failed in clinical use (Poste, 2011). Even large-scale cancer cell line projects, such as the Cancer Cell Line Encyclopedia (CCLE) (Barretina et al., 2012) and the Genomics of Drug Sensitivity in Cancer (GDSC) (Yang et al., 2013) project, do not provide data for cetuximab response. Consequently, the limited amount of cell line and patient data prohibits the use of established statistical methods for biomarker development for cetuximab responsive patient subgroups.

In recent years, several studies showed that mechanistic dynamical models provide an alternative route to biomarker development (Kim and Schoeberl, 2015). Fey et al. (2015) predicted the survival of neuroblastoma patients using a mechanistic model of the c-Jun N-terminal kinase (JNK) pathway. Hass et al. (2017) predicted ligand dependence of solid tumors using a mechanistic multi-pathway model. Frö hlich et al. (2018) showed that large-scale mechanistic models facilitate the integration of large-scale data and enable the derivation and mechanistic interpretation of biomarkers. All these studies employ ordinary differential equation (ODE) models to describe the biochemical reaction networks involved in intracellular signal processing. The models integrate (i) prior knowledge on the pathway topology derived over the last decades and available in databases, such as KEGG (Kanehisa et al., 2010), Reactome (Croft et al., 2011) and BioModels (Li et al., 2010), with (ii) heterogeneous experimental data for the process of interest. The exploitation of prior knowledge constrains the search space and improves in many cases the predictive power. Furthermore, the chosen modeling approach facilitates, unlike most statistical approaches, the mechanistic interpretation of the findings.

In this study, we employed mechanistic mathematical modeling based on ODEs to understand response and resistance mechanisms for cetuximab treatment in gastric cancer cell lines. Building on previously published mechanistic ODE models (Sasagawa et al., 2005; Schö berl et al., 2009; Fujita et al., 2010; Hass et al., 2017; Frö hlich et al., 2018) and published logical models (Flobak et al., 2015), we developed multiple candidate models for the EGFR, the Protein Kinase B (AKT) and the Extracellular-signal Regulated Kinase (ERK) signaling in cetuximab responder and non-responder cell lines. The most appropriate model was selected, calibrated and validated using published and unpublished data. To analyze the dependence of the cellular response on gene expression levels and (somatic) mutations, the resulting model was interrogated using simulation studies and *in silico* knockdown and knockout experiments. This suggests several intervention points and model-based biomarkers.

## 2 Results

### 2.1 Mathematical model of intracellular signaling in cetuximab responder and nonresponder cell lines

We utilized the gastric cancer cell lines MKN1 and Hs746T as a model system to study response and resistance factors of cetuximab treatment. Based on previous results obtained by proliferation and motility analysis, the MKN1 cell line is characterized as a cetuximab responder, while Hs746T is characterized as a non-responder cell line (Kneissl et al., 2012; Keller et al., 2017).

Cetuximab targets the EGFR signaling pathway which regulates growth, survival, proliferation, differentiation (Oda et al., 2005) and motility (Wells, 1999; Keller et al., 2017). Upon ligand binding, EGFR homodimerizes and auto-phosphorylates promoting its catalytic activity. Phosphorylated EGFR (pEGFR) is internalized and degraded or recycled. Membrane-bound and internalized pEGFR catalyzes the activation of the G-protein RAS and the Phosphoinositid-3-Kinase (PI3K) by phosphorylation. While RAS activates the mitogen-activated (MAPK) signaling pathway resulting in the phosphorylation of ERK, PI3K activity results in the phosphorylation of AKT. Phosphorylated ERK (pERK) and phosphorylated AKT (pAKT) are important regulators of DNA transcription.

Cetuximab binds EGFR and blocks the binding of EGF or other EGFR ligands, such as amphiregulin (AREG) and epiregulin (EREG) (Li et al., 2005). This reduces – in the absence of resistance factors – the activity of EGFR and its downstream targets. Known resistance factors to EGFR-targeted therapies in other cancer types include mutations and overexpression of the receptor tyrosine kinases AXL (Hrustanovic et al., 2013) or MET (Zhao et al., 2017).

To quantify differences between the cetuximab responder and non-responder gastric cancer cell lines, we developed a mechanistic model of EGFR signaling and the influence of cetuximab treatment (Figure 1A). The model is based on the available literature, in particular the work of Sasagawa et al. (2005) and Fujita et al. (2010). To limit the model complexity, adaptor proteins and multi-site phosphorylation of EGFR and ERK are disregarded, and the intermediate steps in the MAPK signaling pathway are lumped into a single reaction. We consider wild-type proteins as well as mutant forms, i.e. PI3K p.E545K and MET p.L982 D1028del (den Dunnen et al., 2016). The mutant PI3K expressed in MKN1 cells has an increased catalytic activity compared to wild-type PI3K, resulting in enhanced downstream signaling (Kang et al., 2005). In Hs746T cells MET is mutated and amplified, and assumed to be constitutively active, i.e. independent of its ligand. For detailed information about the model reactions refer to Supplementary Material, Section 3.

**Figure 1:**
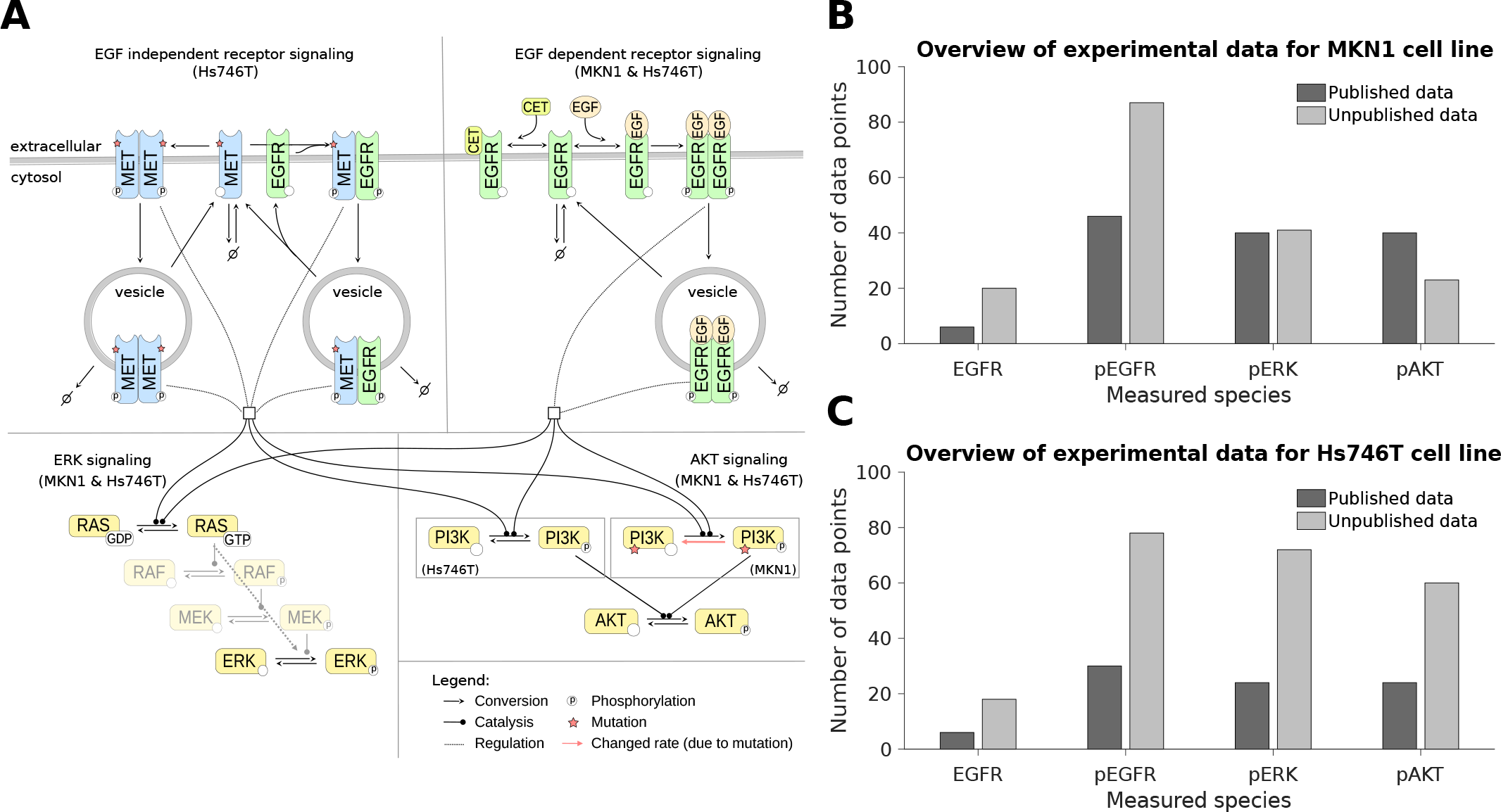
The pathway model and experimental data. (A) Illustration of the mathematical model indicating shared and cell-line specific biochemical species/reactions. (B-C) Overview of the experimental data used for model calibration for MKN1 and Hs746T cell lines. The contribution of literature (published) data and newly collected data (unpublished) is distinguished.

We calibrated the pathway model using a comprehensive dataset obtained using quantitative immunoblotting. The dataset contains time and dose responses for EGFR, pEGFR, pERK and pAKT. In an iterative process, published data (Keller et al., 2017; Kneissl et al., 2012) were complemented by newly collected data to improve reliability of the parameter estimates and model predictions. In total, we considered 303 data points for the MKN1 cell line (Figure 1B) and 312 data points for the Hs746T cell line (Figure 1C).

We determined the maximum likelihood estimates of the model parameters, e.g. rate constants, using multi-start local optimization (Raue et al., 2013, 2015) (see *Materials & Methods*). The estimation was performed individually for MKN1 and Hs746T cells. For MKN1 cells, the optimizer converged reliably. This is reflected in the clear plateau in the waterfall plot (Figure 2A). For the optimal parameters we observed a good agreement of MKN1 data and model fit (Figure 2B-D) (*ρ* = 0.95). For Hs746T cells, the results of the multi-start optimization do not show clear plateaus, suggesting that the objective function possesses a large number of local optima (Figure 3A). But even so, the best fit found provides an accurate description of the data (Figure 3B-D) (*ρ* = 0.91). A potential source of the convergence problems for Hs746T cells is the low signal-to-noise ratio, which is caused by the limited response to EGF and cetuximab treatment (see, e.g., Figure 3C-D) since this is a non-responder cell line. Still, the good agreement of data and model fit confirmed that the proposed pathway model is appropriate.

**Figure 2:**
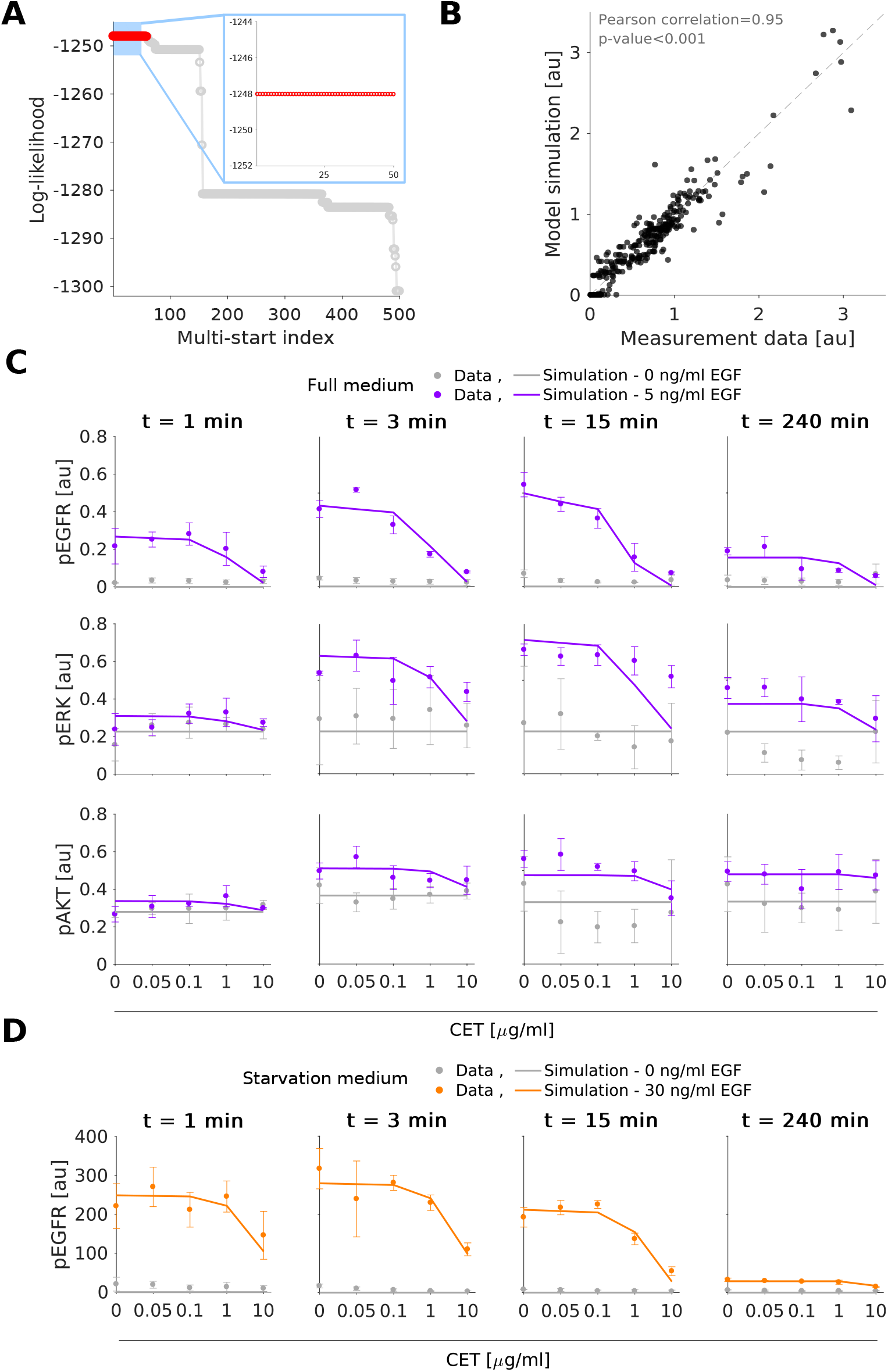
Experimental data and model fit for the gastric cancer cell line MKN1. (A) Waterfall plot for multi-start local optimization. The best 500 out of 1000 runs are depicted from which a magnification of the best 50 multi-starts is indicated by the blue box. Red dots denote the starts converged to the global optimum within a small numerical margin. (B) Scatter plot for the overall agreement of experimental data and model fit. (C-D) Comparison of selected experimental data and model fits. Time and dose response data obtained using immunoblotting indicate the mean and standard deviation of three biological experiments. For visualization in C and D, experimental measurements were scaled to model simulation using the estimated scaling factors to overlay the response to different experimental conditions. Additional data and model fits are provided in Supplementary Material, Section 2.

**Figure 3:**
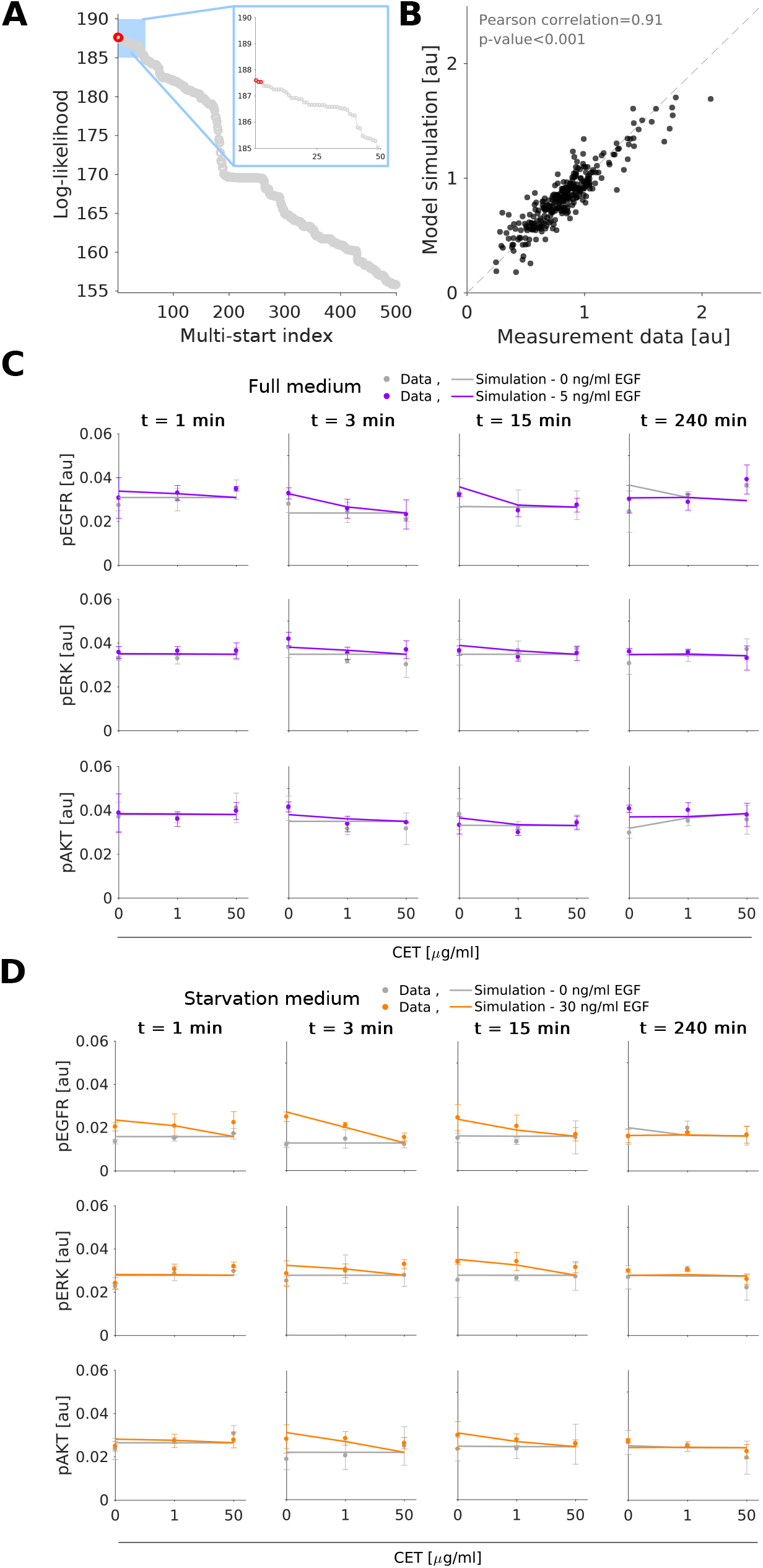
Experimental data and model fit for the gastric cancer cell line Hs746T. (A) Waterfall plots for multi-start local optimization. The best 500 out of 1000 runs are depicted from which a magnification of the best 50 multi-starts is indicated by the blue box. Red dots denote the starts converged to the global optimum within a small numerical margin. (B) Scatter plot for the overall agreement of experimental data and model fits. (C-D) Comparison of selected experimental data and model fit. Time and dose response data obtained using immunoblotting indicate the mean and standard deviation of three biological experiments. For visualization in C and D, experimental measurements were scaled to model simulation using the estimated scaling factors to overlay the response to different experimental conditions. Additional data and model fits are provided in Supplementary Material, Section 3.

### 2.2 Model-based cell line comparison predicts causal differences beyond mutations and expression patterns

The available experimental data (Figures 2 and 3, and Supplementary Material, Section 2) show a pronounced difference in the response of the cell lines to EGF and cetuximab treatment. Potential sources of this behavior are differences in mutation patterns, protein expression levels/abundances and reaction fluxes (due to an influence of latent components between the cell lines). To identify important differences in the reaction fluxes, we compared the parameter estimates obtained for the individual cell lines. These parameter estimates should reflect changes on the biochemical level between the cell lines. In the comparison, we focus on parameters associated with model reactions summarizing multi-step signaling processes. Examples include the indirect activation of ERK by RAS, which is described by a single reaction in the considered pathway model. As the expression levels of the proteins involved in the intermediate reactions – in this case RAF and MEK – can vary, the effective reaction rate constant can easily differ between cell lines. In contrast, for direct reactions, such as the binding of EGF to EGFR, the reaction rate constant should be identical for both cell lines. For details on the classification for the reactions we refer to the Supplementary Material, Section 3.

For the comparison we considered the best 100 parameter vectors obtained by multi-start local optimization for MKN1 and Hs746T cell lines. Statistical testing for cell line specificity of parameters was performed using one-way-ANOVA suggesting significant (*p<* 0.05) differences for 12 out of 20 estimated kinetic rates (Figure 4A). The mapping of the findings on the pathway visualization revealed potential differences in (i) RAS-MAPK signaling, (ii) PI3K-AKT signaling, and (iii) EGFR turnover including internalization, degradation and recycling (Figure 4B). In addition, mutated MET and mutated PI3K can cause differences between cell lines as they are each only present in one of the cell lines.

**Figure 4:**
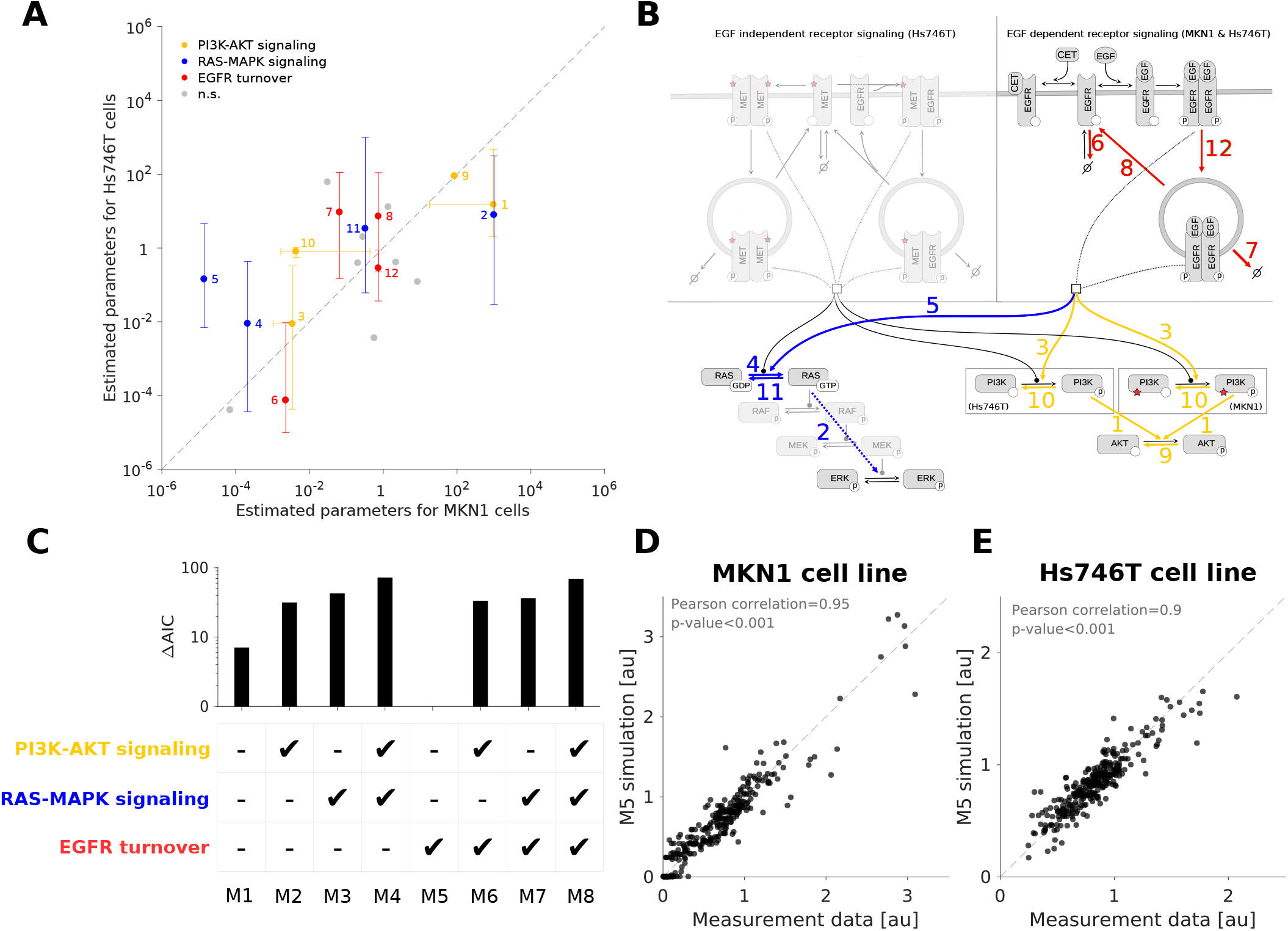
Identification of cell line specific parameters. (A) RAS-MAPK and PI3K-AKT signaling pathways as well as EGFR turnover dynamics as possible cell line specificity candidates. Dots and bars depict the median and one standard deviation (68% percentil interval) of the top 100 log10-transformed parameter estimates obtained by the optimization. Values obtained from MKN1 and Hs746T cells are shown along the X and Y axis, respectively. Coloring indicates significantly different parameter pairs (*p<* 0.05). Gray points indicate non-significantly different and direct reactions (n.s.). (B) Highlighted cell line specificity candidates in the model overview. Coloring and numeric labeling correspond to Figure 4A. (C) Model selection using AIC shows that only changing the receptor turnover dynamics results in the best model. (D) Scatter plot for the overall agreement of experimental data and combined model fit for the MKN1 cell line. Results correspond to the best model (M5). (E) Scatter plot for the overall agreement of experimental data and combined model fit for the Hs746T cell line. Results correspond to the best model (M5). The fits for the selected model (M5) are provided in Figure S1 and Supplementary Material, Section 2.

To assess the relevance of differences in modules (i)-(iii), we fitted the dataset for the MKN1 and Hs746T cell lines simultaneously. Therefore, a collection of mathematical models was developed accounting for cell line specific mutations and protein abundances, as well as differences of parameters belonging to modules (i), (ii) or (iii). All remaining kinetic rates were assumed to be identical for the cell lines. In total, eight different models were constructed accounting for all possible combinations. For each candidate model, a multistart local optimization run was performed. Model selection, using the Akaike Information Criterion (AIC) (Schwarz, 1978), indicated that the model including cell line specific differences only in the EGFR turnover dynamics provides the most appropriate description of the experimental data (Figure 4C). This agrees with previous findings reporting the presence of FBXW7 p.R465C mutation in MKN1 cells, but not in Hs746T cells. FBXW7 ubitiquinates EGFR, leading to changes in the turnover/degradation of EGFR between the cell lines (Heindl et al., 2012). The resulting model provides an accurate description of the experimental data for the responder (Figure 4D) and the non-responder (Figure 4E) cell lines. The subsequent analyzes are based on this model.

### 2.3 Integrated modeling of multiple cell lines yields reliable predictions

The selected pathway model describing the experimental data for the cell lines MKN1 and Hs746T possesses 230 parameters in total. 57 parameters are associated with the reaction kinetics and protein abundances (for wild-type and mutant proteins), and 173 are parameters related to the observations (i.e. scaling constants). To assess the identifiability of the kinetic parameters as well as differences between cell lines and media, we computed the profile likelihoods. We did not calculate the profiles for the observation parameters as they differ between single experiments and are not relevant for the model dynamical response.

Profile likelihoods provide a maximum projection of the likelihood on the individual parameters (*Materials & Methods*). The analysis of the profile likelihoods revealed that 23 out of 57 parameters are practically identifiable, meaning that the 90% confidence intervals are finite (Figure 5A). Most of the identifiable parameters take part in the EGFR dynamics module, involving processes such as internalization, degradation, ligand binding and dimerization. The remaining parameters are practically non-identifiable, meaning that lower and/or upper bounds could not be found for the defined confidence level. In particular, parameters related to MET signaling are practically non-identifiable. This is not unexpected as MMET is not directly observed.

**Figure 5:**
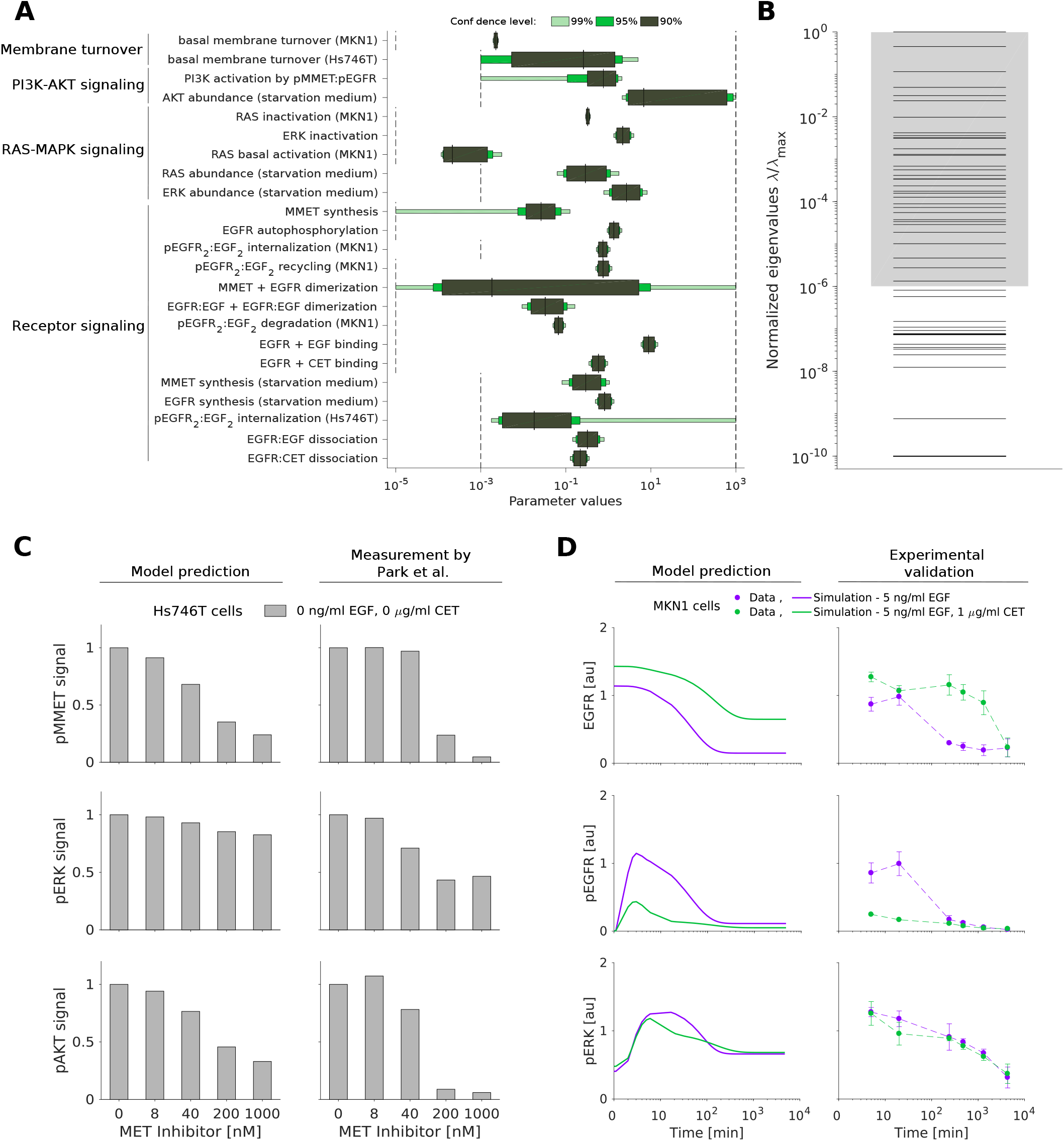
Uncertainty of the parameter estimates does not impair accurate model predictions. (A) Confidence intervals for the identifiable parameters. The confidence intervals corresponding to the confidence levels 90%, 95% and 99% are shown. Parameter bounds used for optimization are indicated in black dashed lines. (B) Eigenvalue spectrum of the FIM for the dynamical parameters. The spectra have each been normalized by their largest eigenvalue (*λ*_max_). Gray shading indicates a spreading of less than six orders of magnitude. (C) *In silico* prediction and experimental validation of MET inhibitor response in Hs746T cells. Validation data for the MET inhibitor KRC-00715 extracted from (Park et al., 2016, Figure 2). The signal was normalized with respect to the untreated condition. (D) Model prediction (left) and experimental validation (right) of long-term response to EGF and the combination of EGF with cetuximab in MKN1 cells.

We complemented the profile likelihood calculation with an evaluation of the Fisher Information Matrix (FIM) at the maximum likelihood estimate. The eigenvalue spectrum of the FIM for the kinetic parameters and initial conditions spans many order of magnitude (Figure 5B) implying sloppiness (Gutenkunst et al., 2007). Interestingly, there are 50 eigenvalues which differ substantially from numerical zero, meaning that the number of constraint directions in parameter space – given by the eigenvectors – is larger than the number of identifiable parameters. This can happen if individual parameters are non-identifiable but functions of several parameters (e.g. sums, differences or ratios) are identifiable. As the detailed analysis of this using profile likelihoods is computationally demanding, the analysis of the FIM provides a first glimpse.

To assess the predictive power of the model despite the sloppy eigenvalue spectrum and the non-identifiable parameters, we simulated the system in additional experimental conditions, i.e. in conditions not used for the fitting. Firstly, we evaluated whether, even though many of the non-identifiabilities are in the MET signaling dynamics, the model can predict published experimental data for the response of Hs746T cells to selected MET inhibitors. We found that the model qualitatively predicts the reduction of pMMET and pAKT levels observed by Park et al. (2016) (Figure 5C). Secondly, we predicted the state of MKN1 cells beyond the first 240 min for which experimental data are available. We found that the model provides reasonable predictions for long-time response of EGFR, pEGFR and pERK (Figure 5D). In summary, this showed that the parameter estimates are reasonable and that the model is predictive.

### 2.4 Mathematical model reliably predicts response and resistance factors

The developed mathematical model provides a screening tool for the rapid assessment of potential response and resistance factors. Here, we demonstrate this by studying the validity of the predictions for several established factors (Figure 6A).

**Figure 6:**
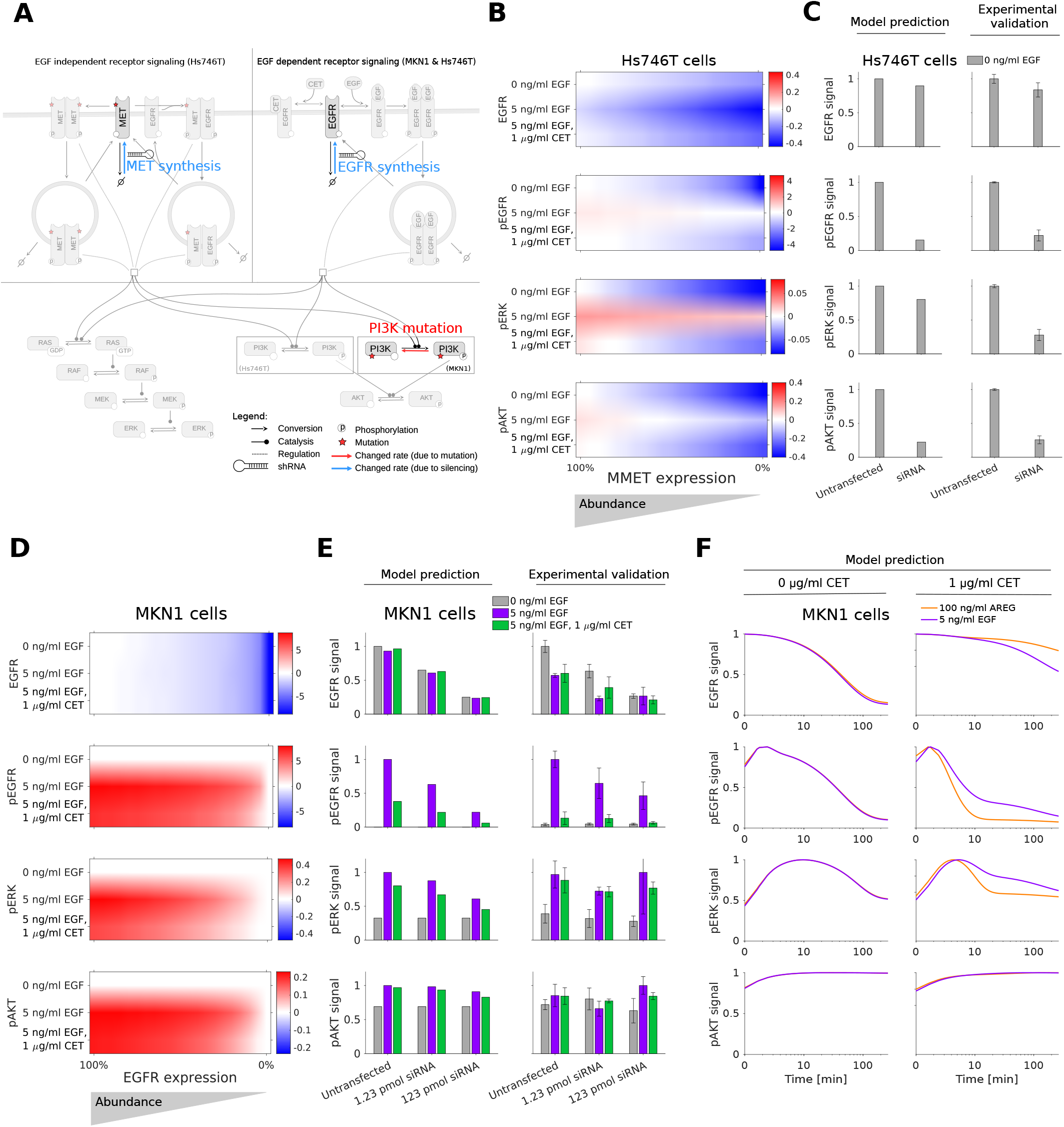
Model prediction of response and resistance factors. (A) Model overview showing changed rates for EGFR and MMET synthesis, and PI3K mutation. (B) *In silico* screening for MMET expression silencing in Hs746T cell line for unstimulated, EGF, and EGF in combination with cetuximab treatment at 3 min. (C) Model prediction and experimental validation at 3 min. (D) *In silico* screening for EGFR expression silencing in MKN1 cell line for unstimulated, EGF, and EGF in combination with cetuximab treatment at 5 min. (E) Model prediction and experimental validation at 5 min. (F) Model prediction of time response of AREG stimulation (black line) shows higher sensitivity to cetuximab treatment compared to EGF (red line) in MKN1 cells. (B-F) The signal is normalized with respect to the maximum activity level for each observed component.

The evident differences between the responder cell line, MKN1, and the non-responder cell line, Hs746T, are the mutation patterns. The experimental data for MKN1 cells showed that PIK3CA p.E545K is not a resistance factor. However, our model predicts the association of PIK3CA overexpression (due to p.E545K mutation) with an insensitivity of pAKT to cetuximab treatment (Figure S2). This is difficult to test experimentally, but consistent with the finding that PIK3CA mutation is a source of acquired cetuximab resistance in metastatic colorectal cancer patients (Xu et al., 2017). The model predictions suggest the same for gastric cancer, although “PIK3CA mutations were not significantly associated with any clinical, epidemiologic, or pathologic characteristic” in gastric cancer patients obtaining non-targeted therapy (Harada et al., 2016).

The MET exon 14 juxtamembrane splice site mutation found in Hs746T cells inhibits the degradation of MET receptor, prolonging its oncogenic activity (Pilotto et al., 2017). MET activation is an established resistance factor for cetuximab treatment in gastric cancer (Kneissl et al., 2012). Indeed, our model predicts that a knockdown of mutant MET reduces Hs746T cell proliferation and survival signaling via ERK and AKT (Figure 6B). To validate the prediction, MET was silenced and quantitative immunoblotting was performed. We found that the model provides accurate quantitative predictions for EGFR, pEGFR, pERK and pAKT (Figure 6C) (Pearson correlation *ρ* = 0.872) although the down-regulation of pERK is slightly underestimated.

Beyond mutations, amplifications and expression changes have been reported as response and resistance factors. In particular the abundance of EGFR has been reported to be associated with cetuximab response in gastric cancer (Zhang et al., 2013; Kneissl et al., 2012). Our model predicts that reducing EGFR expression levels in MKN1 – a cell line overexpressing EGFR – decreases the level of pEGFR, pERK and pAKT (Figure 6D). Interestingly, the effect on downstream signaling was predicted to be relatively small. We tested this prediction by silencing EGFR expression and found a good agreement with experimental data (Figure 6E) (Pearson correlation *ρ* = 0.915). The result implies that the dependence of ERK and AKT activity on EGFR activity is limited.

Beyond the expression of EGFR, the abundance of the EGFR ligand AREG has been shown to correlate positively with cetuximab response (Kneissl et al., 2012). As the biochemical processes underlying AREG- and EGF-induced activation of EGFR signaling are similar (Wilson et al., 2009), we employed the same model for AREG and EGF treatment. Following the literature (Macdonald-Obermann and Pike, 2014), we assumed that AREG has an EGFR affinity about 50 times lower than EGF. We neglected that AREG stimulation leads to EGFR recycling while EGF promotes EGFR degradation (Roepstorff et al., 2009). The resulting model predicted in the absence of MET activation, i.e. in MKN1 cells, that for higher AREG levels cetuximab achieves a higher reduction in EGFR and ERK phosphorylation (Figure 6F). As a consequence of PI3K mutation in MKN1 cells, AKT activation is insensitive to changes in the receptor signal. This model prediction is in line with the published experimental data in (Kneissl et al., 2012).

## 3 Conclusions

Mechanistic ODE models are widely used for the integration of experimental data and the analysis of causal relations. Furthermore, recent studies demonstrate their potential for the identification of biomarkers (Fey et al., 2015; Hass et al., 2017; Frö hlich et al., 2018). To render this approach available for gastric cancer – urgently needed as several large clinical trials failed –, we developed a mechanistic model of signaling in gastric cancer. The model describes the dynamics of the EGFR, ERK and AKT signaling pathways in response to EGF and cetuximab treatment, accounting for mutation patterns and protein expression levels. The proposed model provides a more detailed description than the available logical model (Flobak et al., 2015) by capturing individual biochemical reactions and encoding the wild-type and mutated genes. To the best of our knowledge, the proposed model is the first mechanistic model tailored to gastric cancer signaling.

To assess the predictive power of the model, we performed a broad spectrum of analyzes. First, we used the model to study causal differences between the cetuximab responder cell line, MKN1, and non-responder cell line, Hs746T. The analysis suggested the presence of a MET mutation as well as differences in receptor internalization and recycling as key factors for the response to cetuximab treatment. Second, we validated the predicted response to a MET inhibitor (KRC-00715) as well as to long-term EGF and EGF in combination with cetuximab treatment. Third, we employed the model to predict response and resistance factors. The predicted role of the EGFR abundance as well as MMET were experimentally confirmed by knockdown experiments. The array of successful validations suggests that the model has the potential to be used as a tool for biomarker discovery.

While all tests were positive, the parameters and predictions of the proposed models are subject to uncertainties. Less than 50% of the parameters were practically identifiable, implying that addition of experimental data is required. In particular, measurements of MMET activity would be beneficial, as well as the absolute quantification of expression and phosphorylation levels. Complementary, the model covers only the core pathways and extensions might become necessary. Specifically, additional members of the HER family and additional receptor tyrosine kinases, such as AXL, could be included. A template for the refinement could be provided by the Atlas of Cancer signaling Networks (Kuperstein et al., 2015) or the large-scale mechanistic model we recently introduced (Frö hlich et al., 2018). However, the latter possess a large number of parameters and the acquisition of sufficient experimental data for gastric cancer will be challenging.

In conclusion, the proposed model provides a valuable tool for gastric cancer research. It aggregates the information of a comprehensive list of publications and describes a large set of data points. In the future, it might be used to integrate additional data, e.g. from different cell lines and drugs.

## 4 Materials and methods

### Cell lines and cell culture conditions

For the establishment of the model, the human gastric cancer cell lines Hs746T and MKN1 were used and cultured as reported by Keller et al. (2017).

### Western blot analyzes

Western blot analyzes to set up the model were performed as reported earlier (Kneissl et al., 2012; Heindl et al., 2012; Keller et al., 2017). The cells were treated for indicated times with 0.05, 0.1, 1, 10 or 50 *μ*g/ml cetuximab and/or 5, 30 or 100 ng/ml EGF.

The following antibodies were used: anti-EGFR (Cell signaling Technology (CST), distributed by New England Biolabs in Frankfurt, Germany; #2232; dilution 1:1,000 in 5 % BSA TBS-T), anti-pEGFR Y1068 (Life Technologies, Darmstadt, Germany; #44788G; dilution 1:2,000 in 5 % milk TBS-T), anti-MET (Cell signaling Technology (CST), distributed by New England Biolabs in Frankfurt, Germany; #8198; dilution 1:1,000 in 5 % BSA TBS-T), anti-*α*-tubulin (Sigma-Aldrich, Steinheim, Germany; #T9026; dilution 1:10,000 in 5 % milk TBS-T), anti-*β*-actin (Sigma-Aldrich; A1978; dilution 1:5,000 in 5 % milk TBS-T), anti-mouse IgG (GE Healthcare, distributed by VWR in Ismaning, Germany; #NA931; dilution 1:10,000 in 5 % milk TBS-T) and anti-rabbit IgG (CST; #7074; dilution 1:2,000 in TBS-T).

For quantification, the signals were densitometrically analyzed using ImageJ 1.44p Software (Schneider et al., 2012).

### Knockdown experiments

To generate a transient knockdown of EGFR in the cell line MKN1 and a knockdown of MET in Hs746T cells, siRNA was used. Cells were cultured overnight in rich medium, before the medium was exchanged for antibiotic-free medium. A specific FlexiTube GeneSolution (Qiagen, GS1956 for EGFR, GS4233 for MET) was used, containing four gene specific, preselected siRNAs. The AllStars Negative Control siRNA and AllStars Negative Control siRNA AF488 were used as negative controls. Cells were transfected with Lipofectamine 2000 according to the manufacturer’s instruction. After a transfection time of 4 h (EGFR knockdown in MKN1) or 24 h (MET knockdown in the cell line Hs746T), the medium was replaced with rich medium. The transfection was checked on protein level by western blot analyzes. Therefore, samples were collected 24 hours after the transfection. For analysis of EGFR, MAPK and AKT activation after EGFR knockdown, cells were treated for 5 minutes with 5 ng/ml EGF or the combination of 1 *μ*g/ml cetuximab plus 5 ng/ml EGF.

### Mechanistic modeling

The dynamics of the biochemical reaction networks were modelled using systems of ODEs,

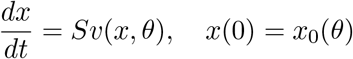

in which *x* denotes the concentration vector of the biochemical species, *S* denotes the stoichiometric matrix, *v*(*x, θ*) denotes the reaction flux vector and *θ* denotes the parameter vector. The parameters are, e.g., rate constants of synthesis or degradation reactions. As the cells start from an unperturbed state, the initial condition *x*_0_(*θ*) is the unperturbed steady state of the ODE.

To infer causal differences between responder and non-responder cell lines, we allowed for differences between the parameters of the cell lines. Following the work of Steiert et al. (2016), we modelled the difference by the additional parameters *ξ*_*i*_,

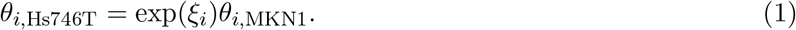

The parameters *ξ*_*i*_ denotes the log-fold change for the *i*-th parameter. If *ξ*_*i*_ is zero, the parameters of the two cell lines are identical. We fixed *ξ*_*i*_ to zero for all parameters which are associate with direct biochemical interactions, e.g. binding rates, which should be conserved between cell lines. Only for expression levels and indirect interactions, i.e. simplification of multi-step processes, we allowed for *ξ*_*i*_ *≠* 0 and therefore estimated along with *θ →* (*θ, ξ*). Similar to the modeling of differences between cell lines, we modelled differences between cell culture media (rich and starvation medium). Here, only differences in the expression levels are allowed. Specific information about parameters are provided in the Supplementary Material, Section 5.

The concentration of biochemical species were linked to the observables *y*(*t*) by observation functions *h*_*j*_,

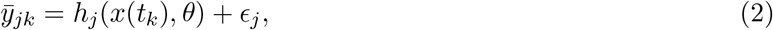

in which *j* indexes the observable and *k* indexes the time point. We assumed independent and additive normally distributed measurement noise 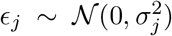. To adapt the scale of different replicates and standard deviation of the experimental data, we employed mixture modeling. The standard deviations were allowed to differ between observables but were assumed to be independent of time and treatment condition for each individual experimental setup (e.g. a set of Western blots). We fixed *σ*^2^ to the estimated standard deviations and therefore reduced the complexity of the optimization problem. Details are provided in the Supplementary Material, Section 1.

### Parameter estimation

We performed maximum likelihood estimation by minimizing the negative log-likelihood function,

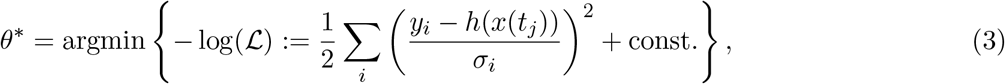

in which *x*(*t*_*j*_) denotes the solution of the ODE model for the parameter vector *θ*.

Numerical optimization was conducted using multi-start local optimization. This approach has been shown to outperform most global optimization methods and achieves a performance comparable with sophisticated hybrid optimization methods (Raue et al., 2013; Villaverde et al., 2018). For each fitting problem, initial parameters were generate using Latin hypercube sampling. The local optimization was performed by trustregion based algorithms implemented in the MATLAB function *lsqnonlin*. For the optimization, parameters were log_10_-transformed to improve numerical properties, e.g. improvement of convexity (Hass et al., 2019) and computational efficiency (Kreutz, 2016; Villaverde et al., 2018). For detailed information about the optimization options refer to Supplementary Material, Section 6.

For the fitting of the individual cell lines, we generated 1000 starting points for local optimization. For the model selection, 100 starting points were used instead. The model selection was based on the AIC values for each model alternative. Following the work of Burnham and Anderson (2002), a difference of 10 was considered to be substantial.

For the assessment of differences between cell lines, we employed an iterative process instead of the regularization approach proposed by Steiert et al. (2016). We also tested the latter, but the optimization did not converge and the penalization did not achieve a clear model reduction.

### Uncertainty analysis

The uncertainty of the model parameters was assessed by analyzing the eigenvalue spectrum of the FIM at the maximum likelihood estimate. The parameter estimation results were considered sloppy if minimum and maximum eigenvalue differed by more than six orders of magnitude (Gutenkunst et al., 2007).

The local analysis provided by the FIM was complemented by computing the profile likelihoods. From the profile likelihoods, we computed the confidence intervals for each individual parameter disregarding scaling constants. Parameters with unbounded confidence intervals for a significance level of 0.1 were considered as practically non-identifiable.

### Software

The model formulation, parameter estimation and uncertainty analysis was performed in the MATLAB toolbox `Data2Dynamics` (https://github.com/Data2Dynamics/d2d, commit 9b1c3556) (Raue et al., 2015). The parameter confidence intervals and the visualization of parameter uncertainties was carried out using the MATLAB toolbox `PESTO` (https://github.com/ICB-DCM/PESTO, commit 8278a1a) (Stapor et al., 2018). For image quantification, we used `ImageJ` software (Schneider et al., 2012).

### Data availability

The experimental data for Hs746T and MKN1 cell lines as well as the complete code – including the toolboxes version – used for the analysis are available at http://doi.org/10.5281/zenodo.2908234. In addition, the SBML model for the biological processes is included in the BioModels database for reusability.

## Acknowledgements

This study was supported by the German Federal Ministry of Education and Research (BMBF) within the framework of the e:Med research and funding concept (SYS-Stomach; Grant No. 01ZX1310A and 01ZX1310B).

## Supplementary Figures

**Figure S1:**
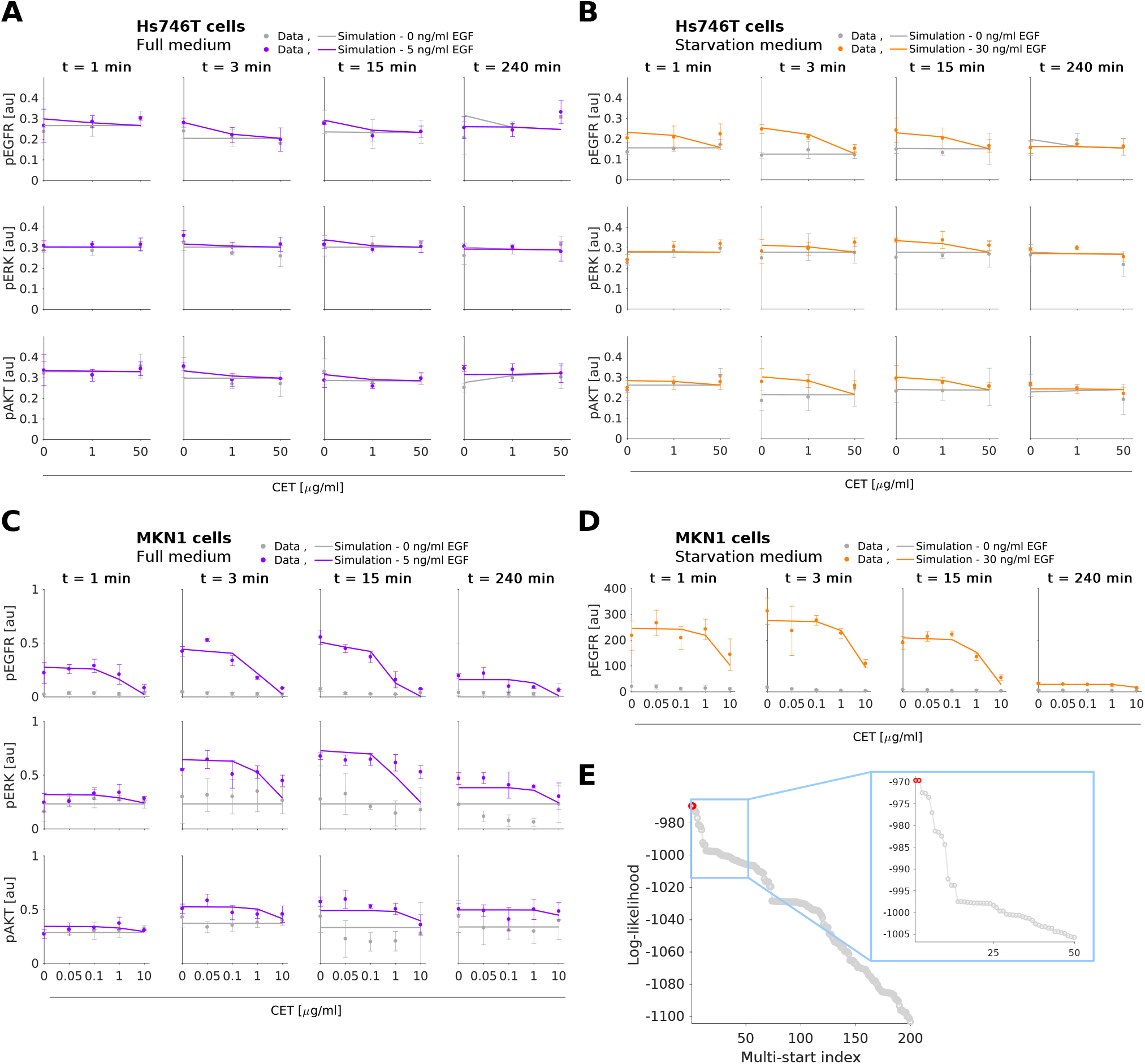
Experimental data and combined model fit for the best model M5. (A-B) Comparison of selected experimental data and model fit for Hs746T cell line. (C-D) Comparison of selected experimental data and model fit for MKN1 cell line. (A-D) Time and dose response data obtained using immunoblotting indicate the mean and standard deviation of three biological experiments. Experimental measurements were scaled to model simulation using the estimated scaling factors. (E) Waterfall plots for multi-start local optimization. The best 200 out of 1000 runs are depicted from which a magnification of the best 50 multi-starts is indicated by the blue box. Red dots denote the starts converged to the global optimum within a small numerical margin. Additional data and model fits are provided in Supplementary Material, Section 4.

**Figure S2:**
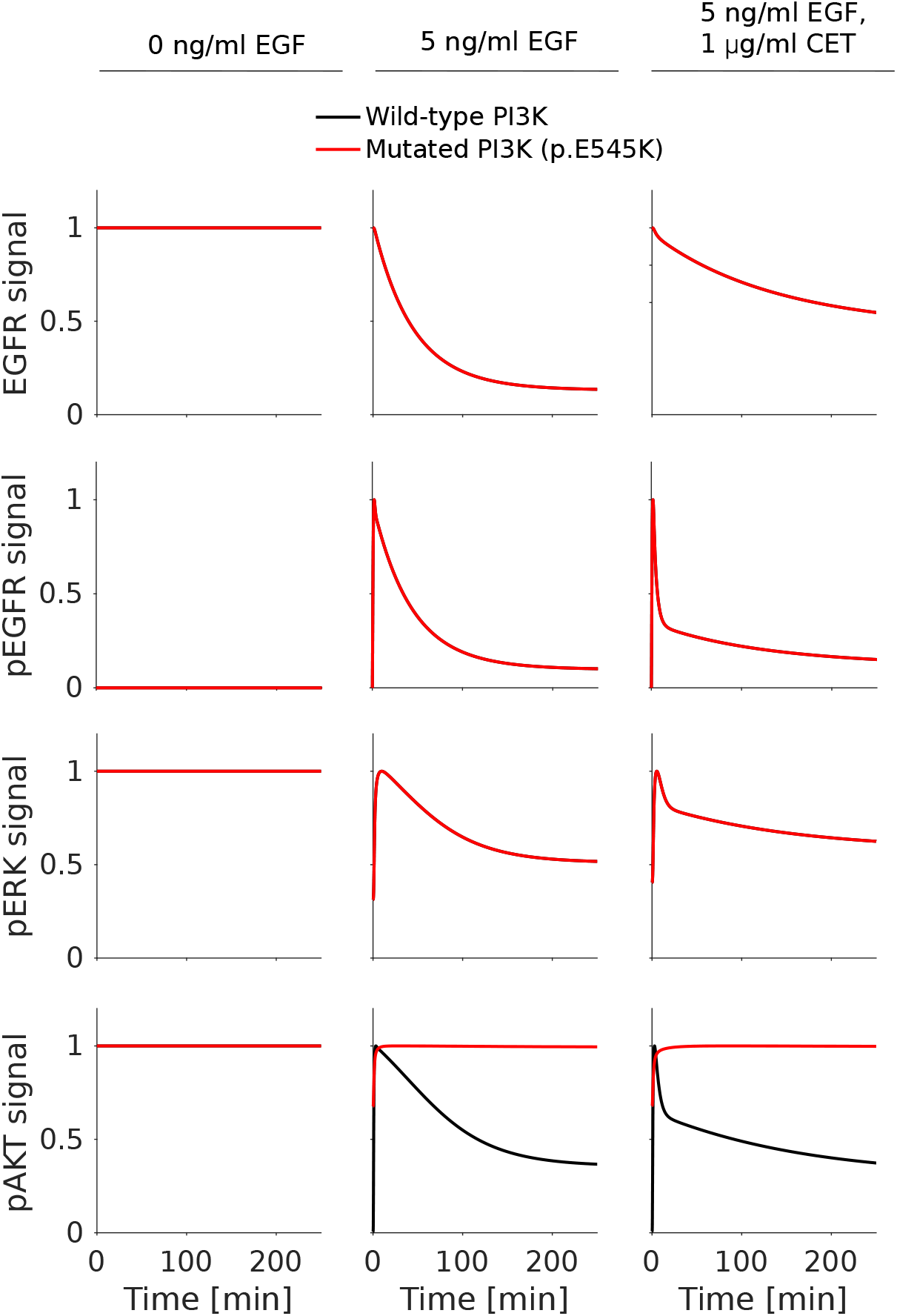
Expression of wild-type PI3K recovers pAKT levels sensitivity to cetuximab treatment in MKN1 cell line. Model prediction of time response of wild-type PI3K (black line) shows a reduction in AKT activity compared to PI3K p.E545K (red line), which remains insensitive. The signal is normalized with respect to the maximum activity level for each observed component.

## Supplementary Material

## 1 Data pre-processing

In this study, we consider a comprehensive collection of Western blot data. As only intensity values on a single gel are comparable, the mapping of the data to the model simulation requires scaling factors (see [Loos et al., 2018] for a detailed discussion). Yet, due to the large number of Western blots, more than one hundred scaling factors would be necessary. This increases the dimensionality of the parameter estimation problem and can cause convergence problems. The problem can be eliminated using novel hierarchical optimization methods [Loos et al., 2018], however, this approach is not supported by `Data2Dynamics` – the modeling toolbox we used for this study.

To reduce the number of scaling parameters, we pre-processed the experimental data by aligning replicates. Replicates are in this case the experimental data for different Western blots capturing the same experimental conditions. Mathematically, we consider the experimental data

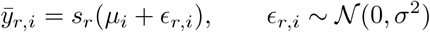

with *r* = 1,*…,R* indexing the replicate and *i* = 1,*…,I* indexing the experimental condition. The experimental condition is a combination of cell line, treatment and time point. The true level of a protein of phospho-protein in experimental condition *i* is denoted by *μ*_*i*_, and corrupted by measurement noise *ϵ*_*r,i*_. The scaling factor associated to different replicates (or Western blots) are denoted by *s*_*r*_. As 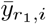 and 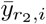 should agree up to measurement noise and scaling factor, we inferred estimates for all but one scaling factor by solving the optimization problem

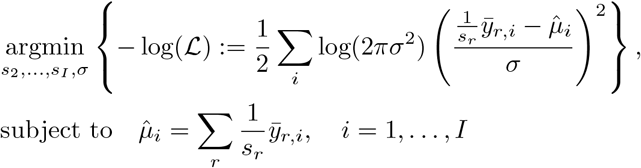

with *s*_1_ = 1 for *r* = 1. The result of the optimization are the pre-processed data 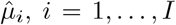 and the estimated standard deviation *σ*. The pre-processed data 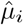 aggregated the information of the replicates by compensating for the individual scales. Without lost of generality, the scale of the first replicate was considered as reference (*s*_1_ = 1), hence, 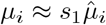 (up to measurement noise).

The pre-processing reduced the number of scaling factors for an experimental dataset from *R* to 1 and renders the optimization problem more tractable. This scaling factor effectively corresponds to the scaling factor for the first replicate *s*_1_.

## 2 Model fits to experimental data

In the main manuscript parts of the large experimental dataset and the corresponding model simulations for the individual fitting of the cell line MKN1 (Figure 2), the individual fitting of the cell line Hs746T (Figure 3), and the combined fitting of MKN1 and Hs746T for the best model (M5) (Figure 5 and Figure S1) are shown. The large datasets were complemented by smaller and more unstructured datasets. These datasets and the corresponding model simulations are depicted in Figures S1, S2, S3 and S4.

**Supplementary Figure S1:**
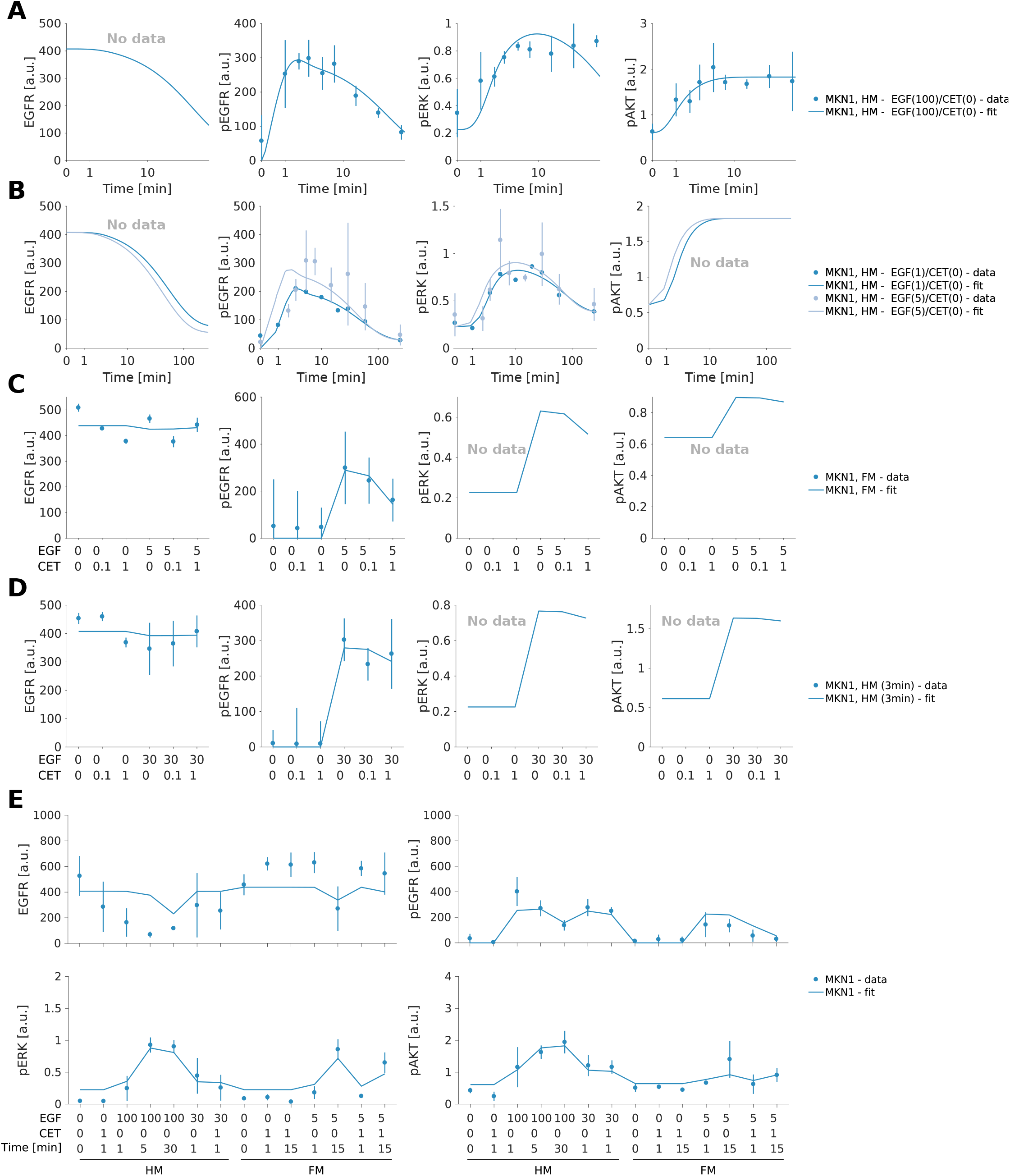
Model-data comparison for the MKN1 cell line, for datasets not depicted in main manuscript Figure 2. (A-B) Time response to different EGF concentrations in starvation culture media (HM). (C) Dose response to EGF and cetuximab stimulation at 3 min in rich culture media (FM). (D) Dose response to EGF and cetuximab stimulation at 3 min in starvation culture media (HM). (E) Dose response to EGF and cetuximab stimulation at 0, 1, 15 and 30 min in full (FM) and starvation culture media (HM). (C-E) Specific EGF and cetuximab concentrations are shown along the X axis.

**Supplementary Figure S2:**
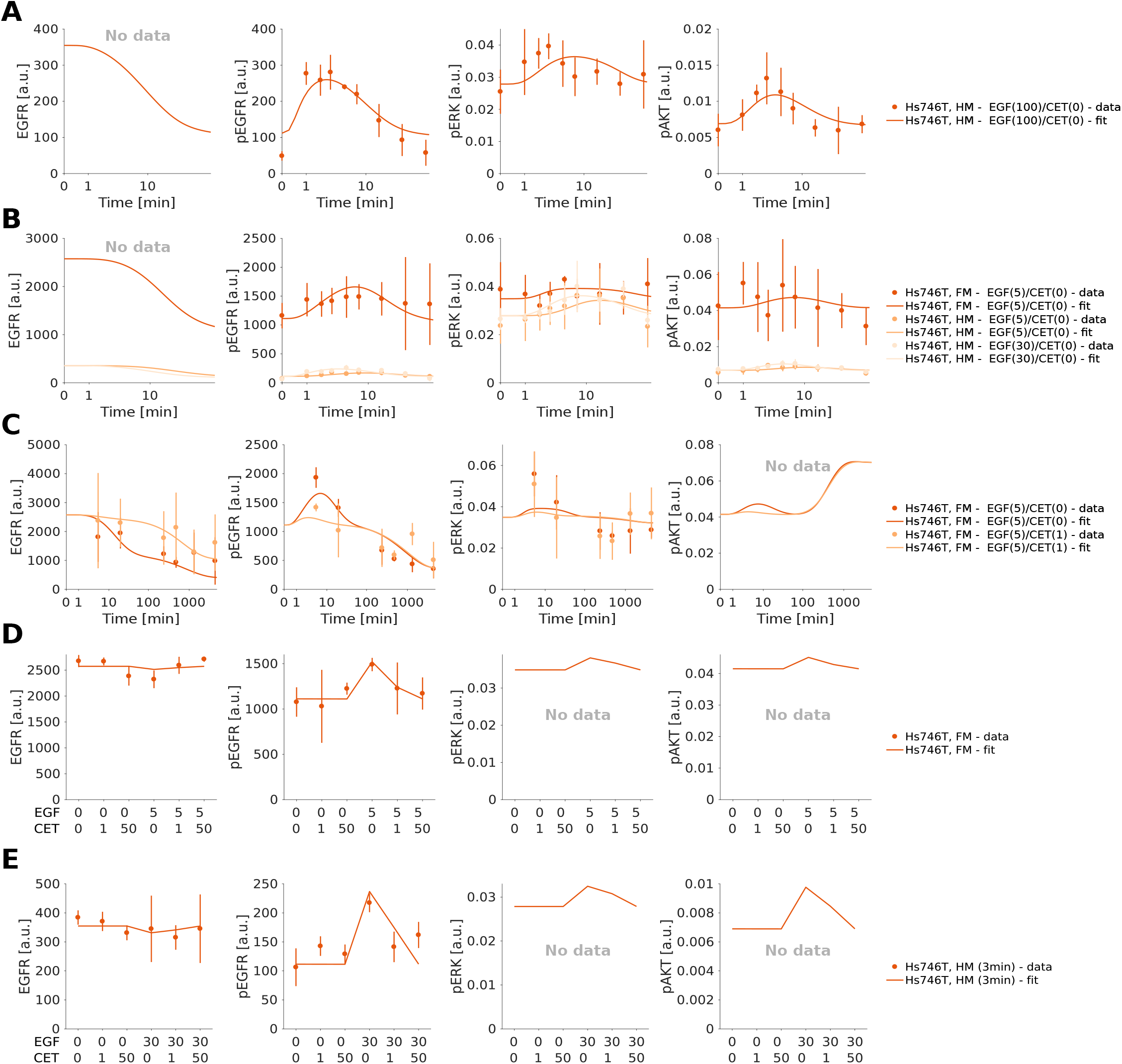
Model-data comparison for the Hs746T cell line, for datasets not depicted in main manuscript Figure 3. (A) Time response to EGF stimulation in starvation culture media (HM). (B) Time response to EGF stimulation in full (FM) and starvation culture media (HM). (C) Time response to EGF and cetuximab stimulation in rich culture media (FM). (D) Dose response to EGF and cetuximab stimulation at 3 min in rich culture media (FM). (E) Dose response to EGF and cetuximab stimulation at 3 min in starvation culture media (HM). (D-E) Specific EGF and cetuximab concentrations are shown along the X axis.

**Supplementary Figure S3:**
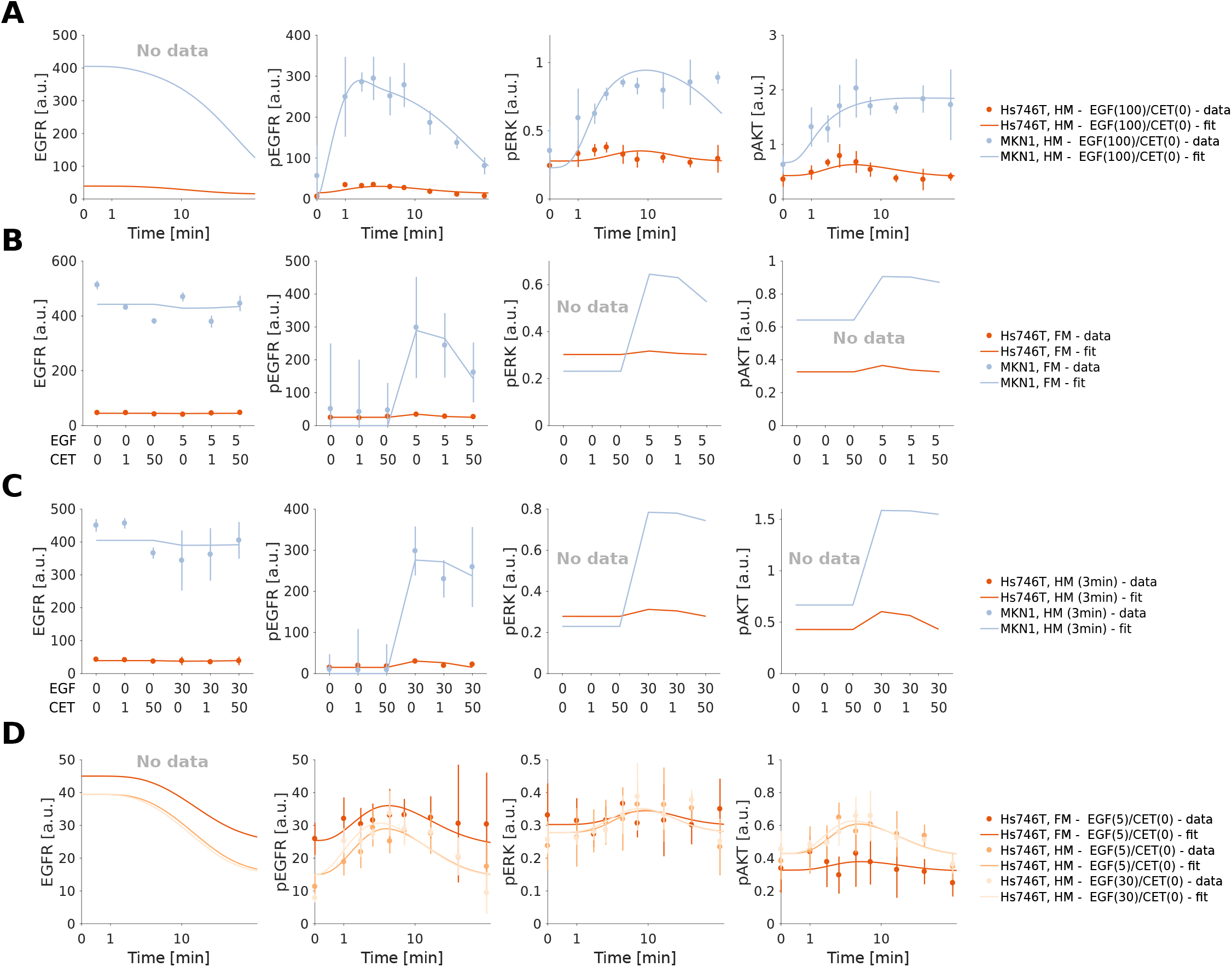
Model-data comparison for the combined fitting of MKN1 and Hs746T cell lines, for datasets not depicted in main manuscript Figure 3 and Figure S1. Model fits for the best model (M5). (A) Time response to EGF stimulation in starvation culture media (HM). (B) Dose response to EGF and cetuximab stimulation at 3 min in rich culture media (FM). (C) Dose response to EGF and cetuximab stimulation at 3 min in starvation culture media (HM). (D) Time response to EGF stimulation of Hs746T cells in full (FM) and starvation culture media (HM). (A-C) Experimental data for both cell lines. (B-C) Specific EGF and cetuximab concentrations are shown along the X axis.

**Supplementary Figure S4:**
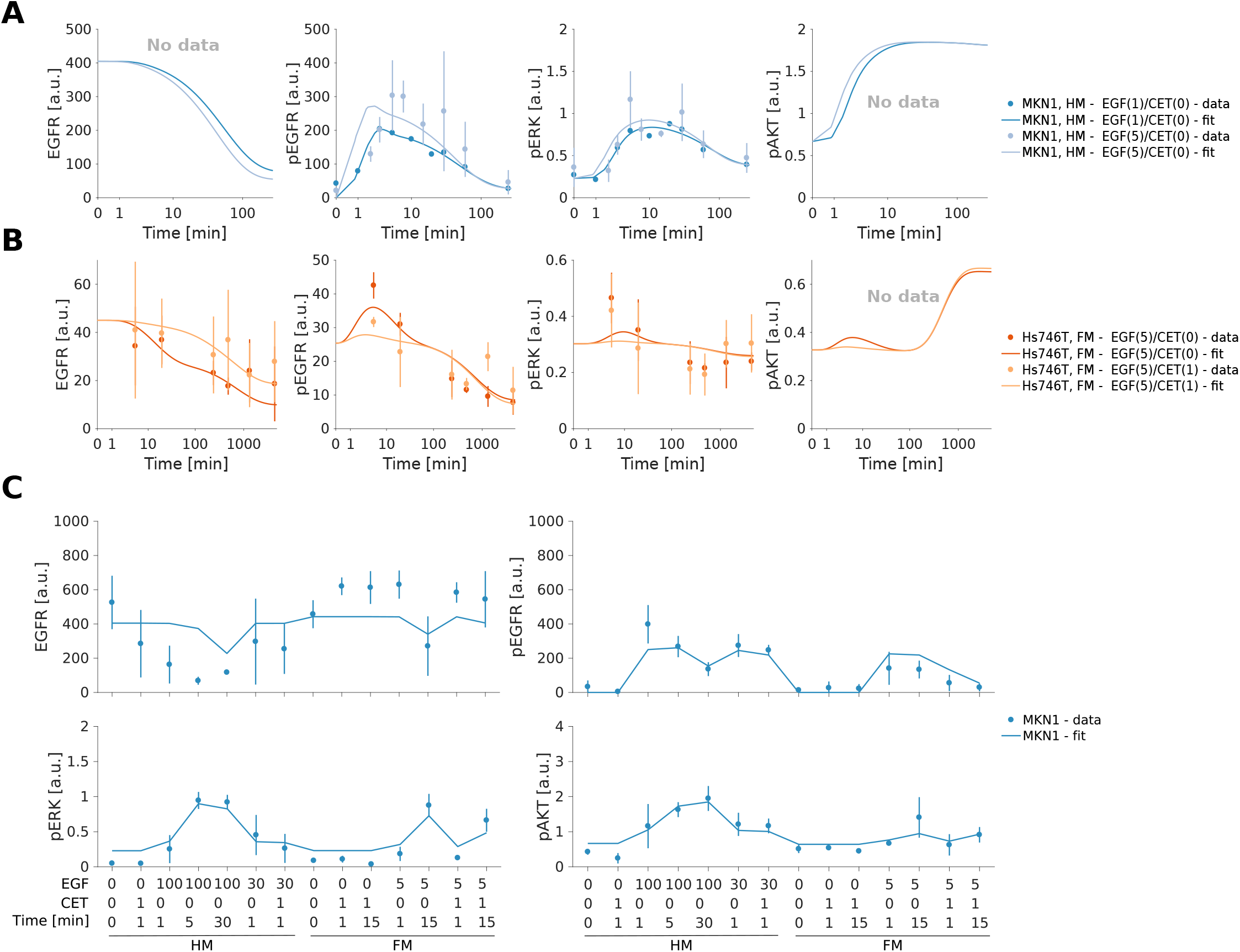
Model-data comparison for the combined fitting of MKN1 and Hs746T cell lines, for datasets not depicted in main manuscript Figure 3 and Figure S1. Model fits for the best model (M5). (A) Time response to EGF and cetuximab stimulation of MKN1 cells in starvation culture media (HM). (B) Time response to EGF and cetuximab stimulation of Hs746T cells in rich culture media (FM). (C) Dose response to EGF and cetuximab stimulation at 0, 1, 15 and 30 min of MKN1 cells in rich (FM) and starvation culture media (HM). Specific EGF and cetuximab concentrations, time points and culture media, are shown along the X axis.

## 3 Model reactions

In the main manuscript a distinction between single reactions describing multi-step signaling processes and describing direct reactions is introduced, which is then used to identify cell line specific reaction rates (Figure 4). In this section, the biochemical reactions implemented in the proposed pathway model are reported. The parts that are exclusive for Hs746T cells are MMET receptor dynamics and EGFR:MMET heterodimer dynamics. Additionally, PI3K is only expressed in Hs746T cells, while MPI3K only in MKN1 cells. The remaining reactions and biochemical species are common for both cell lines. All model reactions follow mass action kinetics. For a more detailed description – including reaction rates – refer to the SBML model.

### EGF receptor dynamics

R1: ∅→ EGFR

R2: EGFR →∅ (*, membrane turnover)

R3: EGFR + CET → EGFR:CET

R4: EGFR:CET → EGFR + CET

R5: EGFR:CET →∅ (*, membrane turnover)

R6: EGFR + EGF → EGFR:EGF

R7: EGFR:EGF → EGFR + EGF

R8: EGFR:EGF + EGFR:EGF → (EGFR:EGF)_2_

R9: (EGFR:EGF)_2_ → (pEGFR:EGF)_2_

R10: (pEGFR:EGF)_2_ → (pEGFR:EGF)_2,endocytic vesicle_ (*)

R11: (pEGFR:EGF)_2,endocytic vesicle_ → EGFR + EGFR (*)

R12: EGFR:EGF →∅ (*, membrane turnover)

R13: (EGFR:EGF)_2_ →∅ (*, membrane turnover)

R14: (pEGFR:EGF)_2,endocytic vesicle_ →∅ (*)

This module consists of 14 reactions describing – among other things – synthesis, ligand and cetuximab (CET) binding, dimerization, internalization and degradation of EGFR. In total 7 out of 14 reactions summarize multi-step processes, denoted by (*). For the dynamics of R2, R5, R12 and R13, we assumed a common kinetic rate describing the membrane turnover. Therefore, from these 7 reactions, 4 different kinetic rates were considered as candidates for cell-line specificity.

### MMET receptor dynamics

R15: ∅→ MMET R16: MMET →∅

R17: MMET + inhibitor → MMET:inhibitor

R18: MMET:inhibitor → MMET + inhibitor

R19: MMET:inhibitor →∅

R20: MMET + MMET → MMET_2_ R21: MMET_2_ → pMMET_2_

R22: pMMET_2_ → pMMET_2,endocytic vesicle_

R23: pMMET_2,endocytic vesicle_ → MMET + MMET

R24: MMET_2_ →∅

R25: pMMET_2,endocytic vesicle_ →∅

R26: MMET + MMET:inhibitor → MMET_2_:inhibitor

R27: MMET:inhibitor + MMET:inhibitor → (MMET:inhibitor)_2_

R28: MMET_2_ + inhibitor → MMET_2_:inhibitor

R29: MMET_2_:inhibitor + inhibitor → (MMET:METinh)_2_

R30: MMET_2_:inhibitor →∅

R31: (MMET:inhibitor)_2_ →∅

This module consists of 16 reactions describing – among other things – synthesis, ligand and MMET inhibitor binding, dimerization, internalization and degradation of MMET. This signaling module is specific for Hs746T cells since MMET is not expressed in MKN1 cells. Consequently, we excluded this group of reactions from the statistical analysis presented in the main manuscript (Figure 4) and, therefore, we did not highlight the different reaction types.

### EGFR:MMET heterodimer dynamics

R32: MMET + EGFR → MMET:EGFR

R33: MMET:EGFR → pMMET:pEGFR

R34: pMMET:pEGFR → pMMET:pEGFR_endocytic vesicle_

R35: pMMET:pEGFR_endocytic vesicle_ → MMET + EGFR R36: MMET:EGFR →∅

R37: pMMET:pEGFR_endocytic vesicle_ →∅

R38: MMET:inhibitor + EGFR → EGFR:MMET:inhibitor

This module consists of 7 reactions describing the dynamics for the EGFR and MMET heterodimers. This signaling module is specific for Hs746T cells since MMET is not expressed in MKN1 cells. Consequently, we excluded this group of reactions from the statistical analysis presented in the main manuscript (Figure 4) and, therefore, we did not highlight the different reaction types.

### RAS-MAPK downstream signaling

R39: RAS-GDP → RAS-GTP (*)

R40: RAS-GDP + (pEGFR:EGF)_2_ → RAS-GTP + (pEGFR:EGF)_2_ (*, EGFR induced activation)

R41: RAS-GDP + (pEGFR:EGF)_2,endocytic vesicle_ → RAS-GTP + (pEGFR:EGF)_2,endocytic vesicle_ (*, EGFR induced activation)

R42: RAS-GDP + pMMET:pEGFR → RAS-GTP + pMMET:pEGFR

R43: RAS-GDP + pMMET:pEGFR_endocytic vesicle_ → RAS-GTP + pMMET:pEGFR_endocytic vesicle_R44: RAS-GDP + pMMET_2_ → RAS-GTP + pMMET_2_

R45: RAS-GDP + pMMET_2,endocytic vesicle_ → RAS-GTP + pMMET_2, docytic vesicle_

R46: RAS-GTP → RAS-GDP (*)

R47: ERK + RAS-GTP → pERK + RAS-GDP (*)

R48: pERK → ERK

This module consists of 10 reactions describing – among other things – basal and receptor induced activation (including EGFR, MMET and their association), and downstream signaling of RAS. We excluded the reactions involving the biochemical species pMMET_2_ and pMMET:pEGFR – also their corresponding internalized forms – as they are only expressed in Hs746T cells. This leads to a total of 5 out of 10 reactions summarizing multi-step processes, denoted by

(*). For the dynamics of R10 and R41, we assumed a common kinetic rate describing the EGFR induced activation of RAS. Therefore, from these 5 reactions, 4 different kinetic rates were considered as candidates for cell-line specificity.

### PI3K- and MPI3K-AKT downstream signaling

R49: PI3K → pPI3K

R50: PI3K + (pEGFR:EGF)_2_ → pPI3K + (pEGFR:EGF)_2_

R51: PI3K + (pEGFR:EGF)_2,endocytic vesicle_ → pPI3K + (pEGFR:EGF)_2,endocytic vesicle_

R52: PI3K + pMMET:pEGFR → pPI3K + pMMET:pEGFR

R53: PI3K + pMMET:pEGFR_endocytic vesicle_ → pPI3K + pMMET:pEGFR_endocytic vesicle_

R54: PI3K + pMMET_2_ → pPI3K + pMMET_2_

R55: PI3K + pMMET_2,endocytic vesicle_ → pPI3K + pMMET_2,endocytic vesicle_

R56: pPI3K → PI3K (*, †) R57: MPI3K → pMPI3K

R58: MPI3K + (pEGFR:EGF)_2_ → pMPI3K + (pEGFR:EGF)_2_

R59: MPI3K + (pEGFR:EGF)_2,endocytic vesicle_ → pMPI3K + (pEGFR:EGF)_2,endocytic vesicle_

R60: MPI3K + pMMET:pEGFR → pMPI3K + pMMET:pEGFR

R61: MPI3K + pMMET:pEGFR_endocytic vesicle_ → pMPI3K + pMMET:pEGFR_endocytic vesicle_

R62: MPI3K + pMMET_2_ → pMPI3K + pMMET_2_

R63: MPI3K + pMMET_2,endocytic vesicle_ → pMPI3K + pMMET_2,endocytic vesicle_

R64: pMPI3K → MPI3K (*, †)

R65: AKT + pPI3K → pAKT + pPI3K (*, PI3K/MPI3K induced activation)

R66: AKT + pMPI3K → pAKT + pMPI3K (*, PI3K/MPI3K induced activation)

R67: pAKT → AKT (*)

This module consists of 19 reactions describing – among other things – basal and receptor induced activation, and downstream signaling of PI3K – expressed in Hs746T cells – and MPI3K – expressed in MKN1 cells. All kinetic rates defined for PI3K and MPI3K are common between the cell lines, except for the inactivation rate – due to PI3K mutation in MKN1 cells – indicated by (*†*). We excluded the reactions involving the biochemical species pMMET_2_ and pMMET:pEGFR – also their corresponding internalized forms – as they are only expressed in Hs746T cells. This leads to a total of 4 out of 19 reactions summarizing multi-step processes, denoted by (*). We assumed a common kinetic rate describing the PI3K (and MPI3K) induced activation of AKT. Therefore, from these 4 reactions, 3 different kinetic rates were considered as candidates for cell-line specificity.

## 4 Candidate reactions for cell line specificity

In the main manuscript the identification of cell line specific reaction rates is shown (Figure 4). Here, a detailed description of these reaction rates is provided in Supplementary Table S1. For simplicity, in the following table we show the biochemical reactions in which the kinetic rates is involved. Note that the same kinetic rate can appear in different reactions, e.g., rate for basal membrane turnover in R2, R5, R12 and R13. For a more detailed description refer to the SBML model.

## 5 Model and model parameters

The model accounts of 27 biochemical species. Protein abundances for RAS, ERK, PI3K, MPI3K and AKT were assumed to be conserved. This results in an ODE model with 22 state variables. As the absolute abundances of the conserved species are unknown, we consider as state the relative abundance with the concentration in MKN1 cells and rich culture media as reference point for RAS, ERK, MPI3K and AKT. As PI3K is not represented in MKN1 cells, we used as reference point the concentration in Hs746T cells and rich culture media. The changes in expression levels and synthesis between the different culture media and cell lines were estimated as proposed by Steiert et al. [2016].

As initial conditions for the simulations of time and dose response data, we employed the steady state of the unperturbed system. This steady state is specific to the combination of cell line and culture medium.

We screened the literature for the kinetic rates considered in the model. Using the obtained information, we defined prior distributions for individual rate constants. We considered normal distributions of the log-transformed paper values. The normal priors were centered around, i.e. mean, the corresponding literature values with a standard deviation of 0.2. The values are listed in Supplementary Table S2.

**Supplementary Table S1:** Reaction rate candidates for cell-line specificity. The following kinetic rates were identified as significantly different between the cell lines by the ANOVA statistical test. All kinetic rates belonging to the same module were simultaneously tested (not individually) resulting in 8 different model candidates.

| Module | Reaction | Description |
| --- | --- | --- |
| EGFR turnover | R10: $(\text{pEGFR:EGF})_2 \rightarrow (\text{pEGFR:EGF})_{2,\text{endocytic vesicle}}$ | Endocytosis of pEGFR |
| | R14: $(\text{pEGFR:EGF})_{2,\text{endocytic vesicle}} \rightarrow \emptyset$ | Lysosomal degradation of endocytosed pEGFR |
| | R2: $\text{EGFR} \rightarrow \emptyset$ | Basal membrane turnover |
| | R5: $\text{EGFR:CET} \rightarrow \emptyset$ | |
| | R12: $\text{EGFR:EGF} \rightarrow \emptyset$ | |
| | R13: $(\text{EGFR:EGF})_2 \rightarrow \emptyset$ | |
| | R11: $(\text{pEGFR:EGF})_{2,\text{endocytic vesicle}} \rightarrow \text{EGFR} + \text{EGFR}$ | EGFR recycling endosomes |
| RAS-MAPK signaling | R39: $\text{RAS-GDP} \rightarrow \text{RAS-GTP}$ | Basal activation of RAS |
| | R40: $\text{RAS-GDP} + (\text{pEGFR:EGF})_2 \rightarrow \text{RAS-GTP} + (\text{pEGFR:EGF})_2$ | EGFR dependent activation of RAS |
| | R41: $\text{RAS-GDP} + (\text{pEGFR:EGF})_{2,\text{endocytic vesicle}} \rightarrow \text{RAS-GTP} + (\text{pEGFR:EGF})_{2,\text{endocytic vesicle}}$ | |
| | R46: $\text{RAS-GTP} \rightarrow \text{RAS-GDP}$ | Basal GTPase activity of RAS |
| | R47: $\text{ERK} + \text{RAS-GTP} \rightarrow \text{pERK} + \text{RAS-GDP}$ | RAS dependent activation of MAPK |
| PI3K-AKT signaling | R65: $\text{AKT} + \text{pPI3K} \rightarrow \text{pAKT} + \text{pPI3K}$ | PI3K and MPI3K dependent activation of AKT |
| | R66: $\text{AKT} + \text{pMPI3K} \rightarrow \text{pAKT} + \text{pMPI3K}$ | |
| | R67: $\text{pAKT} \rightarrow \text{AKT}$ | Basal dephosphorylation of pAKT |
| | R56: $\text{pPI3K} \rightarrow \text{PI3K}$ | Basal dephosphorylation of pPI3K and pMPI3K |
| | R64: $\text{pMPI3K} \rightarrow \text{MPI3K}$ | |

## 6 Optimization settings

For the parameter optimization of the MKN1 model, Hs746T model and combined model, the MATLAB functions *lsqnonlin* and *fmincon* were considered. For the MKN1 model and the combined model, *lsqnonlin* achieved the better optimization results. For the Hs746T model, *fmincon* achieved the better results. For each model, we used the best results across the optimization algorithms. Based on the optimizer performance observed for the combined model, the model selection was performed using *lsqnonlin*.

For all the calibrated models – regardless of the optimizer – the following optimizer tolerances we

- maximum number of iterations allowed (*MaxIter*) was set to 10^4^,
- maximum number of function evaluations allowed (*MaxFunEvals*) was set to 10^4^,
- termination tolerance on the function value (*TolFun*) was set to 10^-9^, and
- termination tolerance on *x* (*TolX*) was set to 10^-10^,

and the following ODE solver tolerances were used

- relative tolerance (*rtol*) was set to 10^-8^,
- absolute tolerance (*atol*) was set to 10^-8^, and
- maximum number of integration steps (*maxsteps*) was set to 10^6^.

The remaining settings for optimizer and ODE solver were set to the default values provided by `Data2Dynamics.`

**Supplementary Table S2:** Literature values used as prior information in the parameter estimation. Prior mean values were converted to 10-logarithmic scale and used for the parameterization of the model. Note that the same kinetic rate can appear in different reactions, e.g., receptor endocytosis in R10, R22 and R34.

| Reaction | Value | Publication |
| --- | --- | --- |
| R6, R7: $\text{EGFR} + \text{EGF} \rightleftharpoons \text{EGFR:EGF}$ | $\log_{10}(2)$ | Klein et al. [2004], page 1 |
| R3, R4: $\text{EGFR} + \text{CET} \rightleftharpoons \text{EGFR:CET}$ | $\log_{10}(0.39)$ | [Kim and Grothey, 2008], page 2 |
| R10: $(\text{pEGFR:EGF})_2 \rightarrow (\text{pEGFR:EGF})_{2,\text{endocytic vesicle}}$<br>R22: $\text{pMMET}_2 \rightarrow \text{pMMET}_{2,\text{endocytic vesicle}}$<br>R34: $\text{pMMET:pEGFR} \rightarrow \text{pMMET:pEGFR}_{\text{endocytic vesicle}}$ | $\log_{10}(0.25)$ | [Sorkin and Duex, 2010], page 19 |
| R67: $\text{pAKT} \rightarrow \emptyset$ | $\log_{10}(1.5 \times 60)$ | [Schöberl et al., 2009], kinetic rate $k_f83$ |

